# Stretching combined with repetitive small length changes of the plantar flexor muscles enhances their passive extensibility for longer duration than conventional static stretching, while not compromising strength

**DOI:** 10.1101/333864

**Authors:** Naoki Ikeda, Takayuki Inami, Yasuo Kawakami

## Abstract

Static stretching increases flexibility but can decrease muscle strength, and the method to avoid the latter has been longed for. In this study, a novel stretching modality was developed that provides repetitive small length changes to the plantar flexor muscles undergoing passive static stretching (“local vibration stretching,”). We investigated the effects of local vibration stretching on muscle strength, flexibility and its persistence. Plantar flexion strength and maximal ankle joint dorsiflexion angle (dorsiflexion range of motion) were measured for 10 healthy young males before (pre) and immediately after (post) three types of stretching: static stretching, local vibration stretching at 15 Hz, and no intervention (control). The dorsiflexion range of motion was measured also at 15, 30, and 60 min post-stretching. Elongation of the medial gastrocnemius and Achilles tendon was determined by ultrasonography. Plantar flexion strength significantly decreased by 4.3 ± 3.5 % in static stretching but not in local vibration stretching. The dorsiflexion range of motion significantly increased both in static stretching (7.2 ± 8.1 %) and local vibration stretching (11.2 ± 14.6 %) which was accompanied by a significantly larger muscle elongation but not tendon elongation. Elevated dorsiflexion range of motion was maintained until 30 min after the local vibration stretching while it returned to baseline level (pre-intervention) in 15 min after the static stretching. All variables remained unchanged in the control condition. In conclusion, local vibration stretching improves extensibility of the muscle belly without decreasing strength, and the increased flexibility is retained longer than static stretching.

## Introduction

Modalities to improve flexibility (joint range of motion) are roughly divided into static stretching (SS) in which muscles are stretched while holding the joint at a fixed angle, and dynamic stretching (DS) where muscles experience dynamic stretch–shortening within the maximal range of motion. Although flexibility can be significantly improved after static stretching, muscle strength and functions are often attenuated [1,2]. The latter outcome has been attributed to the reduction of the neural drive [3,4,5] and a decrease in the muscle-tendon unit (MTU) stiffness [3,6,7]. In contrast, DS provides a smaller effect on flexibility than static stretching but it does not sizably attenuate muscle functions [8,9,10]. Possible factors involved in DS include dynamic stretching and shortening of actively contracting muscles, which might be responsible for the smaller negative effect of DS on muscle functions. This can be due to a retained neural drive and/or a negligible change in MTU stiffness after DS [10,11].

Previous animal [12] as well as human [13] studies showed a decrease in MTU or muscle stiffness that underwent cyclic stretch–shortening cycles passively. There is a possibility that such a modality leads to further improvement of flexibility than DS, while posing no negative effect on muscle functions unlike SS. Performing DS passively and cyclically with a small range of joint angle (5° for instance), can be such a modality. When this maneuver is performed at a relatively high frequency, it can be regarded as “vibration”. Vibration stimuli to the body or muscles provide a positive effect on muscle functions, e.g., muscle force enhancement [14,15,16], and also improves flexibility [14,17]. Thus, a modality that conditions the MTU by SS combined with dynamic length changes by vibration will be effective in improving muscle stiffness while retaining muscle functions. To the best of our knowledge, no study has ever tried such a stretching maneuver.

In the present study we developed a novel stretching technique which employs the feature of DS (in the form of vibration) added onto SS, for the purpose to take advantages of both SS and DS. We named this technique as “local vibration stretching (LVS)”. Attempts to apply vibration stimuli to the target muscles undergoing static stretching have been performed [18,19]. In those studies, vibration stimuli were applied using a vibrator on the muscle belly [18] or a whole-body vibration device [19]. The vibration amplitude was very small in those approaches with its direction not specified along the target muscles. The present concept of LVS is essentially different from those approaches because the vibration stimulus with a sizable amplitude (15mm: ankle joint angle change of approximately 5°) is provided to the major plantar flexors, thereby applying repetitive small length changes of MTU like DS under passive stretching.

The altered flexibility is reported to persist for 10 - 30 min [2,10]. after static stretching, and at least 10 min after the DS [11], and at least 15 min after the vibration stimulus [17]. Combination of these interventions may further elongate their after-effects, but this idea has not been tested.

In the present study, we conducted the experiment on the MTU of the lower leg for the purpose of verifying the effects of LVS on muscle strength, flexibility, and persistence of altered flexibility. It was hypothesized that LVS does not decrease muscle strength while improving and maintaining flexibility similarly to static stretching.

## Materials and methods

### Participants

The participants were 10 recreationally active males without apparent neurological, orthopedic, or neuromuscular problems of their lower legs (age, 22 ± 2 years; body height, 1.70 ± 0.06 m; body weight, 64.3 ± 8.9 kg; mean ± SD). The details and purpose of this study as well as the risk associated with participating in this study was explained to each participant in advance, before obtaining consensus for participation. This study was approved by the Ethics Review Committee on Human Research of Waseda University and performed in accordance with the Declaration of Helsinki.

### Study Design

The present study was aimed to clarify the effects of LVS on muscle strength, flexibility, and persistence of altered flexibility, and comparing those effects with SS. The right ankle joint was tested for all participants, in the following three conditions: SS intervention, LVS intervention, and no stretching (control). The participants were tested under these conditions in a random order, with an interval of 3 days or longer, after measuring the maximal voluntary plantar flexion torque to determine the basis for dorsiflexion range of motion (DFROM) measurement as described later.

### Measurement of maximal voluntary muscle strength

Isometric maximal voluntary plantar flexion torque was measured by using an isokinetic dynamometer (VTF-002, VINE, Japan), with the knee fully extended in a sitting position, and the ankle secured to a foot plate of the dynamometer at 0° (anatomical position). The signal obtained from the dynamometer was amplified by an amplifier (DPM-711B, Kyowa, Japan), then digitally converted at 1000 Hz through an A/D converter (PowerLab/16SP, ADInstruments, Australia), fed into a personal computer (FMV Lifebook, Fujitsu, Japan) and recorded by using a software (LabChart7, ADInstruments, Australia). Before measuring maximal voluntary plantar flexion torque twice, the participants were instructed to warm-up, producing force below the maximal strength for several times. The peak torque was analyzed per measurement, and the third measurement was performed when the values differed by 5% in two measurements. The higher value in two measurements was taken as maximal voluntary plantar flexion torque. Maximal voluntary plantar flexion torque was measured before and immediately after the intervention.

### Measurement of dorsiflexion range of motion

Flexibility was measured as DFROM by using the same isokinetic dynamometer used in maximal voluntary plantar flexion torque measurement (Fig 1). The ankle joint was dorsiflexed at 5°/s starting from 30° plantar flexion, until the passive resistive torque corresponding to 20% of the pre-measured maximal voluntary plantar flexion torque was reached when the DFROM was measured. The participants were instructed to relax during the measurements without resisting to the passive dorsiflexion. The right foot region including the distal part of the lower leg was videotaped (Exilim, Casio, Japan) at 30 Hz to obtain the joint angle. For this purpose, reflective markers were attached onto the following four landmarks: the upper and lower side of the foot plate, medial malleolus, and tibia (the middle point of the line between the estimated center of the knee joint and the medial malleolus). In the recorded image, the two-dimensional coordinates of those markers were obtained by using a software (FrameDIAS4, DKH, Japan), and the angle between the vectors parallel to the foot plate and the line from the medial malleolus to the tibia was defined as the ankle joint angle, being 0° at the neutral position and positive values for dorsiflexion. The video was synchronized with other data using a synchronizer (PH-100, DKH, Japan). The measurement was repeated twice, and the higher value was adopted. DFROM was measured before, immediately after the intervention (POST), and at 15min (POST 15), 30min (POST 30), and 60min (POST 60) post-stretching to examine the persistence of altered flexibility.

**Fig 1.**
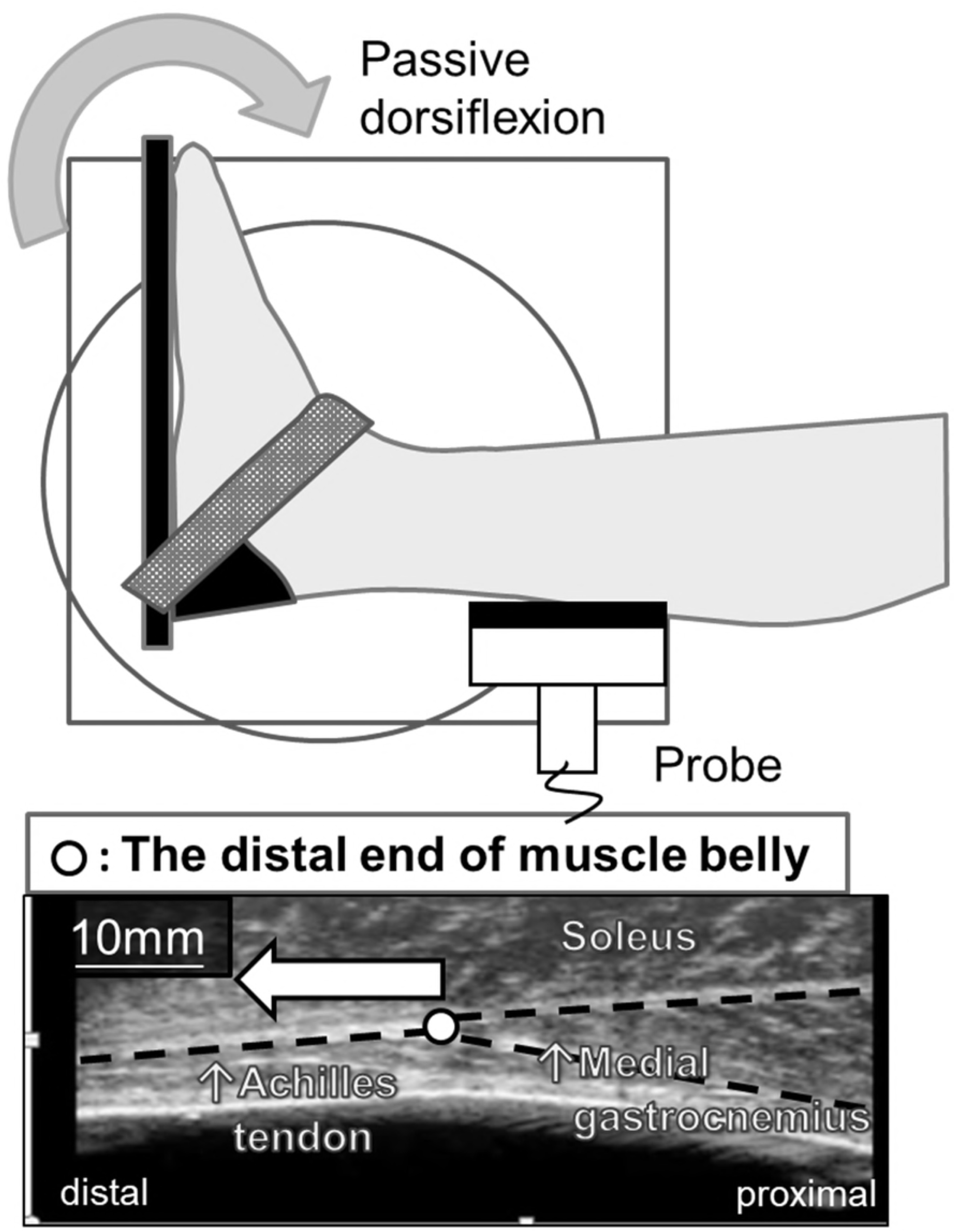
Illustration of DFROM measurement and muscle lengthening.

### Measurement of muscle and tendon elongations

During DFROM measurement pre- and post-intervention, the proximo-distal movement of the distal end of muscle belly of the medial gastrocnemius was recorded by using an ultrasonic apparatus (SSD-6500, ALOKA, Japan, connected to a video recorder GV-HD700, SONY, Japan operating at 30 Hz) to represent muscle elongation [21] (Fig 1). The ultrasonic probe (frequency: 7.5MHz; scan width: 60 mm; UST-5712, Hitachi Aloka Medical, Japan) was fixed onto the skin with a double-sided adhesive tape above the distal end of muscle belly of the medial gastrocnemius. The videotaped ultrasound images were later analyzed using a software (FrameDIAS4, DKH, Japan) to measure the muscle elongation from the position of 30° plantar flexion up to DFROM.

The changes in the length of the medial gastrocnemius–Achilles tendon MTU from 30° plantar flexion to DFROM were estimated by using the following formula [20],

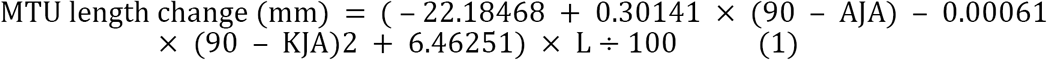

(AJA: ankle joint angle (90° = anatomical position), KJA: knee joint angle (0° = full extension position), L: lower leg length (linear distance [mm] from the popliteal fossa to the lateral malleolus of the ankle joint)

The difference between the change in MTU length and muscle elongation was defined as tendon elongation [21]. The analysis was repeated twice per measurement, and the mean value was adopted. The coefficient of variation of muscle elongation was 0.88% in two measurements on the same participant.

### Stretching protocol

For both SS and LVS, an isokinetic dynamometer (CON-TREX, CMV, Switzerland) with a dynamic stretching device (JM-25, TOPRUN, Japan) mounted on the foot plate, was used. The posture of the participants was the same as that of the DFROM measurement. Based on a previous study [22], the stretching duration was 15min in total for both SS and LVS (15 sets of 1-min stretching, with an interval of 30s). The dorsiflexion angle for SS was set at the same angle as in the maximal joint angle during DFROM measurement at pre-intervention. For LVS, a dynamic stretching device plantar- and dorsiflexed the ankle by approximately 5° (Fig 2) at 15 Hz for 1min, around the same position as in SS. The selection of vibration frequency of 15 Hz was to avoid mechanical stress to the muscle and the feeling of discomfort and pain that were brought about at higher vibration frequencies in a preliminary experiment. The ankle joint angle was positioned at 0° during the interval between sets of LVS. In control, the participants were instructed to sit at rest on the dynamometer for approximately 25 min, while maintaining the right knee and ankle joints lightly flexed and plantar flexed to avoid stretching of plantar flexors.

**Fig 2.**
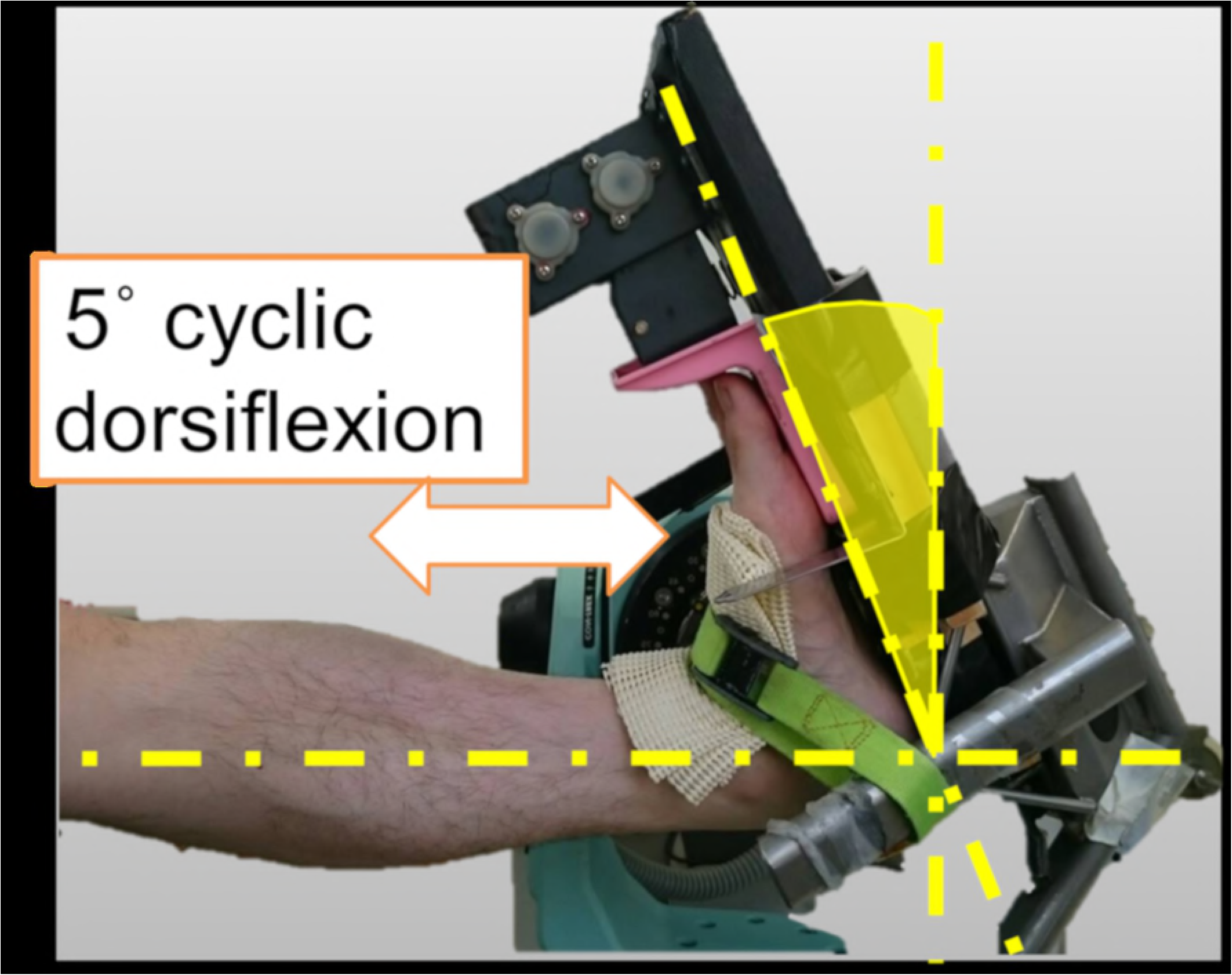
A picture showing implementation of local vibration stretching.

### Statistical analysis

All data are presented in means ± SD. For the values of maximal voluntary plantar flexion torque, DFROM, and muscle and tendon elongations before (pre) and after (post) intervention, two-way repeated measures analysis of variance (ANOVA) was performed on the stretching conditions (control, SS, and LVS) ×; time (pre and immediately post) (SPSS 12.0J, SPSS Japan, Japan). When an interaction or a main effect for time was observed, a paired t-test was performed in each condition. The time course changes in flexibility were compared for each condition. The relative changes in maximal voluntary plantar flexion torque, DFROM, muscle, and tendon elongations before and immediately after intervention were examined for statistical differences using one-way repeated measures ANOVA. When the F value was significant, a Tukey multiple comparison test was performed. For SS and LVS conditions, DFROM values at POST, POST 15, POST 30, and POST 60 were normalized to the pre-measurement values. A two-way repeated ANOVA was performed on the stretching conditions (SS and LVS) ×; time (POST, POST 15, POST 30, and POST 60). When a significant interaction or a main effect for time was observed, a one-way repeated measures ANOVA with least significant difference (LSD) post-hoc test was performed in each condition. Significant differences among pre-measurement values in different conditions were assessed by one-way repeated measures ANOVA for all parameters. The level of statistical significance was set at *p* < 0.05.

## Results

A significant interaction between the conditions and time was observed in maximal voluntary plantar flexion torque, and maximal voluntary plantar flexion torque significantly decreased in SS after intervention, whereas no change was observed in control or LVS (Fig 3). The relative change of maximal voluntary plantar flexion torque after intervention in SS (−4.3 ± 3.5%) was significantly different from that in control (0.0 ± 2.9%), whereas no significant difference was observed between LVS (−1.6 ± 3.9%) and control.

**Fig 3.**
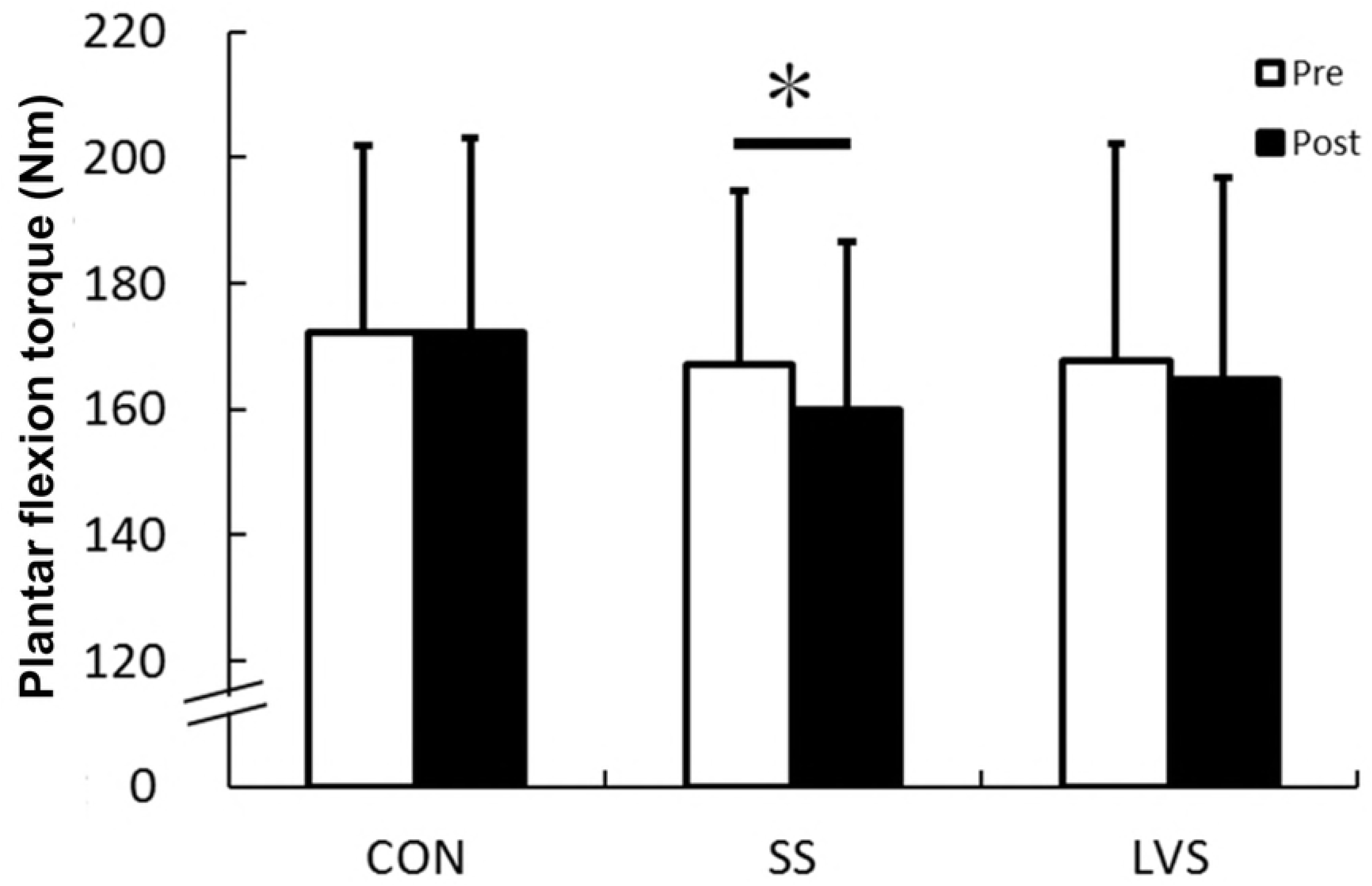
Changes in maximal voluntary isometric plantar flexion torque in each condition. CON: control; SS: static stretching; LVS: local vibration stretching. ^*^: significantly changed compared with pre-intervention (*p* < 0.05). Values are means ±SD

The main effect for time was observed in DFROM, and it significantly increased both in SS and LVS but not in control (Fig 4). The relative change of DFROM after intervention in LVS (11.2 ± 14.6%) was significantly higher than that in control (−0.7 ± 4.0%), whereas no significant difference was observed between LVS and SS (7.2 ± 8.1%), and between SS and control.

**Fig 4.**
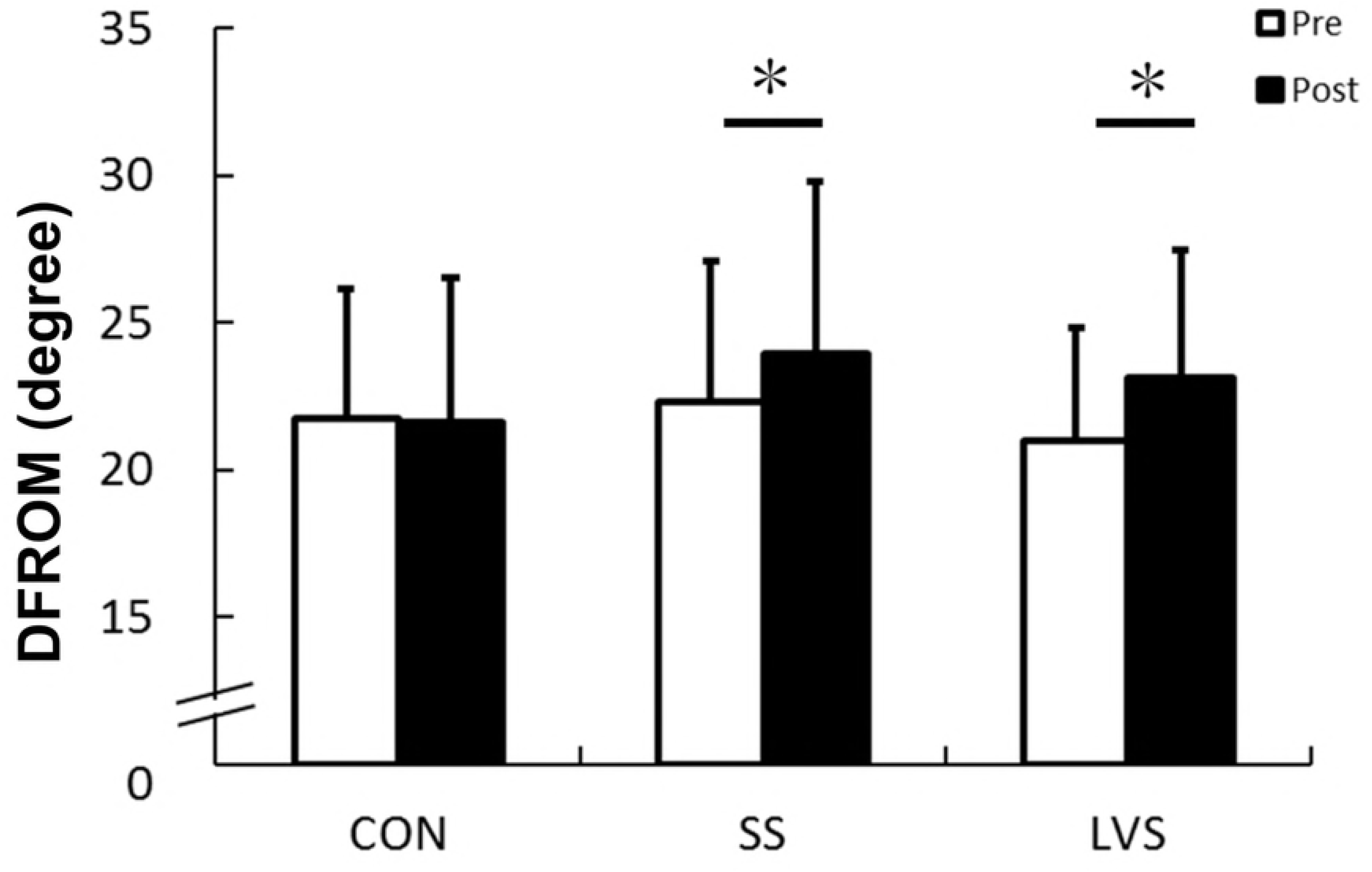
Changes in the dorsiflexion range of motion in each condition. CON: control; SS: static stretching; LVS: local vibration stretching. ^*^: significantly changed compared with pre-intervention (*p* < 0.05). Values are means ±SD

The interaction was observed between conditions and time in muscle elongation, and it significantly increased both in SS and LVS, whereas no change was observed in control (Fig 5). The relative change of muscle elongation after intervention in LVS (8.5 ± 10.2%) was significantly higher than that in control (−0.8 ± 3.2%), whereas no significant difference was observed between LVS and SS (5.8 ± 6.2%), and between SS and control. As for the tendon elongation, the main effect for time was not significant, and no change in any conditions was observed (Fig 6).

**Fig 5.**
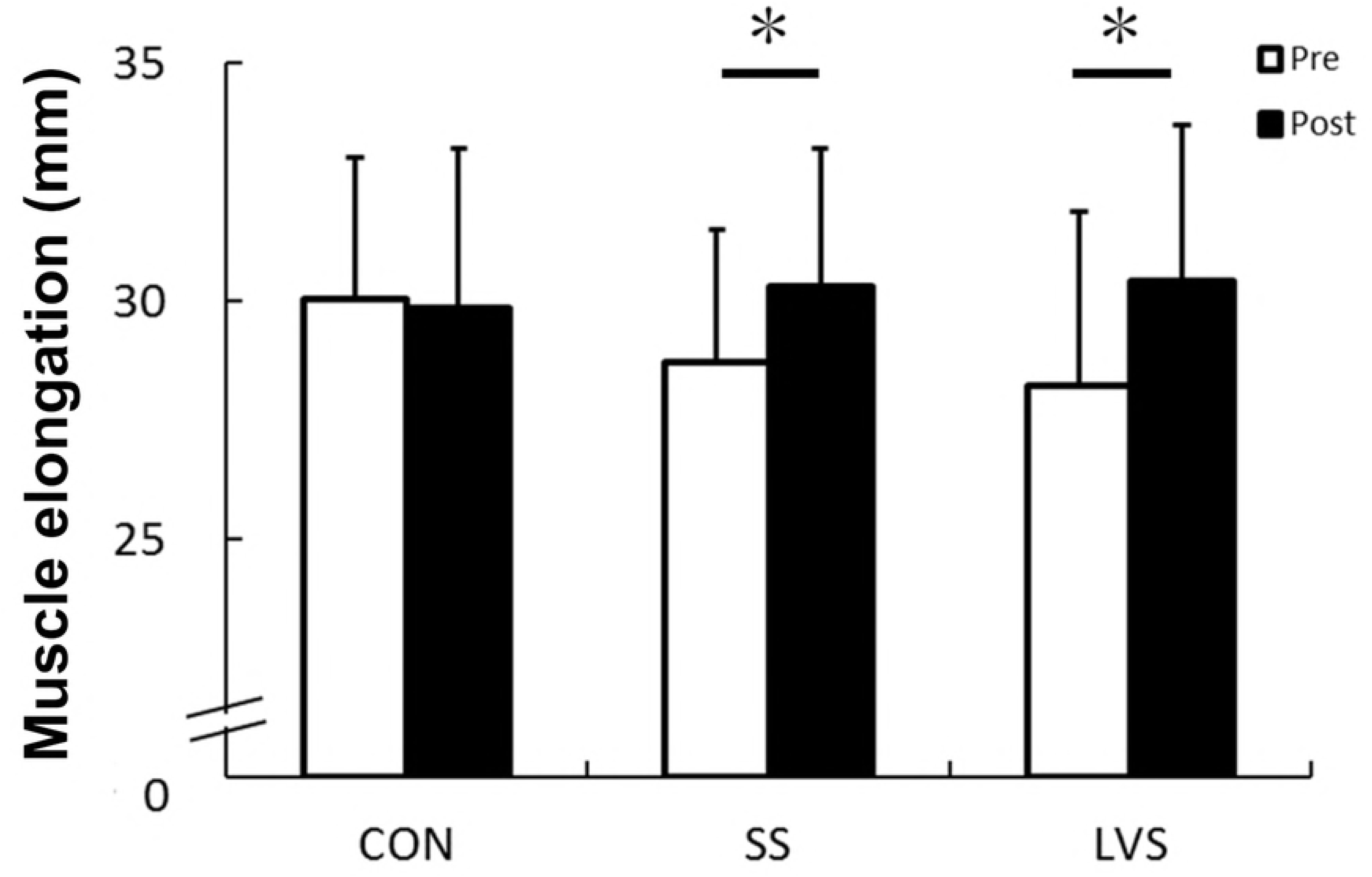
Changes in muscle elongation in each condition. CON: control; SS: static stretching; LVS: local vibration stretching. ^*^: significantly changed compared with pre-intervention (*p* < 0.05). Values are means ±SD (n = 9)

**Fig 6.**
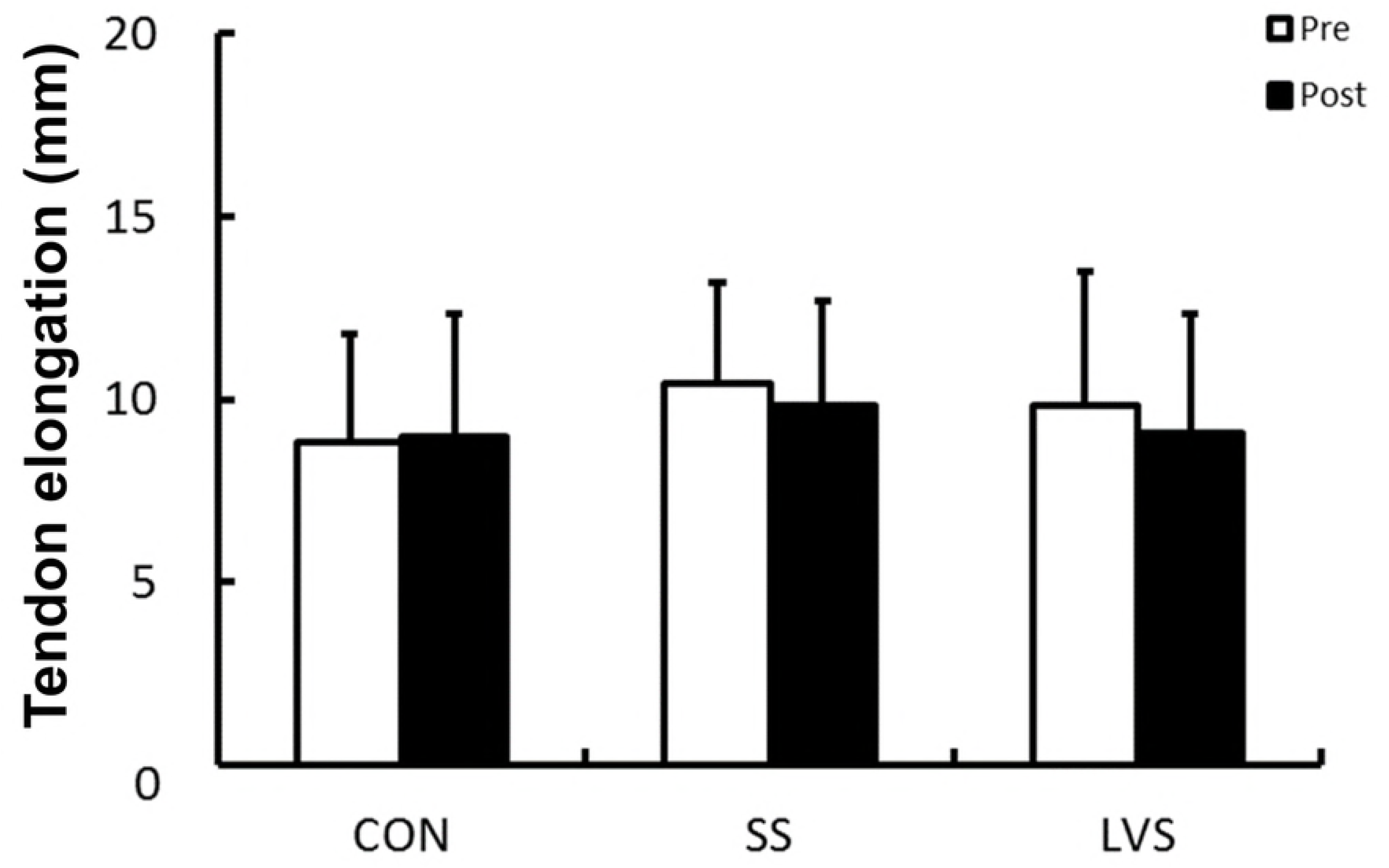
Changes in Achilles tendon elongation in each condition. CON: control; SS: static stretching; LVS: local vibration stretching. No significant difference was observed before and after intervention the conditions (*p* > 0.05). Values are means ±SD (n = 9)

The main effect for time was observed in the change of DFROM relative to the pre-intervention value: in SS, it was significantly smaller at POST15, POST30 and POST60 compared with POST while in LVS, it was significantly smaller at POST60 compared with POST, POST15 and POST30 (Table 1).

**Table 1:** Time course of dorsiflexion range of motion in static stretching and local vibration stretching conditions.

|  | Static stretching (%) | Local vibration stretching (%) |
| --- | --- | --- |
| Immediately post-stretching | 7.2 ± 8.1 | 11.2 ± 14.6 |
| 15 min post-stretching | 0.4 ± 7.9 * | 10.7 ± 11.1 |
| 30 min post-stretching | 0.2 ± 10.6 * | 5.5 ± 13.5 |
| 60 min post-stretching | -1.0 ± 9.4 * | -2.0 ± 12.2 * † ‡ |

\*: significantly different compared with immediately post-stretching ( $p < 0.05$ ). †: significantly different compared with 15 min post-stretching ( $p < 0.05$ ). ‡: significantly different compared with 30 min post-stretching ( $p < 0.05$ ). Values at each time point are the relative changes from the pre-intervention values. Values are means ±SD.

## Discussion

This study revealed the following effects of LVS that muscle strength was not compromised unlike SS, while DFROM was improved to a similar extent to SS. In addition, it was shown that the enhanced DFROM by LVS persisted longer than SS.

Previous studies reported muscle functions (e.g. muscle strength and power) declined after SS [1,4,10]. Our results for SS coincided with these studies. In contrast, maximal voluntary plantar flexion torque was unchanged in LVS, and the relative change of maximal voluntary plantar flexion torque in LVS was not different from that in control, unlike SS. Thus, LVS did not decrease muscle strength. SS is reported to decrease the neuromuscular activity during force production [4,10], but LVS is assumed not to reduce neuromuscular activity, because maximal voluntary plantar flexion torque was unchanged. In contrast, muscle strength/power can be improved after DS [8,10], and this has been attributed to 1) conditioning effect through lengthening and shortening of MTU, and 2) active force production during DS, although their relative contributions are unknown. LVS provided passive and repetitive small length changes to the MTU undergoing SS without active force production, and the muscle strength was not improved. This result suggests that the above factor 2) is likely to be a dominant trigger for improvement of the muscle strength by DS. Although muscle strength transiently increases after being vibrated at about 30Hz [16], this finding is not directly compared to the present study because vibration modalities are completely different as mentioned above.

DFROM significantly and similarly increased in SS and LVS. This suggests that DFROM was increased by LVS with a SS-like effect on the plantar flexor MTU by dorsiflexing the ankle into the final ROM similarly to SS. An increase in muscle elongation due to SS has been thought to be caused by a decrease in passive muscle stiffness and changes in neurophysiological properties including lowered stretch-reflex sensitivity [2,6,23,24]. Muscle elongation was comparable for LVS and SS, suggesting that the muscle was similarly affected by these two interventions, but there also was a tendency of the former (9%) being larger than the latter (6%). This could be explained by a greater decrease in muscle stiffness in LVS compared with SS resulting from passive and cyclic stretch–shortening cycle (as passive DS) and vibration stimuli1 [2,25,26].

No change was observed in tendon elongation in SS or LVS. The load-elongation relationship of the tendinous tissue is divided into the toe region (larger and nonlinear elongation to a smaller tensile force) and the linear region (stiffer and linear elongation-tension relation) [27,28]. DFROM measurement in both SS and LVS was performed at the intensity corresponding to 20%MVC, which may have been within the toe region of the tendon force-length relationship where the effect of stretching on the tendon is not influential [4,29].

The improved flexibility subsided after 30 min in LVS, while within 15 min in SS. Previous studies reported persistence of altered flexibility being 10 min [7] or 30 min [2] after SS, 10 min after DS [11], and 15 min after vibration stimulus alone [17]. The persistence of altered flexibility in this study differs from these reports. This may be because of the differences in methodology, e.g., duration and intensity of interventions. Neurophysiological changes caused by SS can affect the persistence of altered flexibility for ∼2 min, and changes in mechanical properties of muscular and tendinous tissues can keep altered flexibility for 8 min∼ [30]. It appears therefore, that the persisting effect of LVS on flexibility is due to changes in muscle and tendon mechanical properties. Although further studies are required to clarify this mechanism, the present study indicated that LVS enhanced DFROM and it persisted longer than SS.

In the present study, muscle activities were not measured; hence the extent of neuromuscular activity during measurement and stretching is unknown and the above related arguments remains largely speculative. The effects of LVS on neuromuscular activity are worth investigating in future studies. LVS developed in this study clearly differs from SS in that it provides repetitive small length changes to MTU longitudinally (unlike conventional segmental or whole-body vibration), and also differs from typical DS in terms of the lack of active force production; thus, LVS is a novel stretching technique. Examination of factors not dealt with in the present study, e.g., combinations of different stroke lengths and vibration frequencies, might lead to development of more effective application of LVS on flexibility and exercise performance improvement.

## Conclusion

This study revealed the following three effects of a newly developed stretching technique (local vibration stretching: LVS): 1) muscle strength is not compromised unlike SS, 2) DFROM increases to a similar extent as SS, and 3) enhanced DFROM subsided after 30 min in LVS, while it persists longer than in SS.

## Acknowledgments

We sincerely thank Kohei Akase (former undergraduate student at the School of Sport Science, Waseda University) for his cooperation. This study was supported by the JSPS KAKENHI (Grant Number 16H01870).

## Supporting information

**S1 Raw Data.** Changes in strength, dorsiflexion range of motion (DFROM), time course of DFROM, muscle and tendon elongation of each participant in each condition.

